# Characterizing interactions in the nuclear pore complex transporter using novel site-specific deuteration and SANS

**DOI:** 10.1101/2024.06.20.599953

**Authors:** Arpita Tawani, Shuo Qian, Samuel E. Sparks, Piakulla Sashi, Sean Cahill, Michael P. Rout, David Cowburn

## Abstract

We describe an unprecedented SANS/solvent matched experiment with the aim of providing a meso-scale dynamic structural description of fuzzy complexes such as those formed by nuclear transport factors as they ferry cargo across the nuclear pore complex, whose description is normally limited by the large number of states and rapid time scales of interconversion. These contain repeat short linear interaction domains which are common features of many intrinsically disordered functional proteins – e.g., transcriptional regulators, and RNA-interaction domains associated with liquid-liquid phase separated non-membranous organelles. The novel approach uses site-specific deuteration, SANS, and model fitting to provide changes to average spatial distributions between interacting domains of FG Nups upon binding to the nuclear transport factor NTF2. The results support the fully disordered nature of phenylalanyl-glycyl repeats within FG Nups in their interactions with the nuclear transport factor in vitro, as well as the absence of significant inter-aromatic contacts, or of interchain linkage in complexes.

## Introduction

A significant number of molecular cellular events involve dynamic complexes with functional involvement of intrinsically disordered proteins and their interaction partners on multiple time scales including sub micro-seconds (1). Examples include transcription complexes(2), assembly and disassembly of nucleosomes (3, 4), mechano-transduction (5) and regulators of post translational modification (6). An extreme example is the functional role of the intrinsically disordered phenylalanyl-glycyl rich nucleoporin segments (FG Nups) in the nuclear pore complex (7) where disorder plays a thermodynamic and kinetic role in facilitated versus passive transport (8).

The complexity of describing such systems generally is well recognized (9) and regarded as a major challenge to understanding the molecular basis of such interactions and their biological effects (10).

Small angle scattering (11) (SAS) methods provide substantial information about the size and shape of such systems (12), and neutron scattering (SANS) provides the unique ability to scatter selectively by specific selective deuteration of components (13). Selective deuteration of synthetic polymers e.g. (14) is established to determine their properties, and the potential is recognized for site-(15) or sequence-selective deuteration of proteins (16).

Selective transport of macromolecules between the nucleus and cytoplasm is mediated by nuclear pore complexes (NPCs), embedded in the nuclear envelope. The central channel of NPCs is occupied by a cloud of intrinsically disordered regions containing multiple Phe-Gly (FG) repeats, anchored in the channel wall; the proteins responsible are termed FG Nups. Cargoes are trafficked through this cloud by association with nuclear transport factors (NTRs), whose specific interaction with the repeat’s multiple Phe residues gives them access to this dense environment that otherwise excludes non-specific macromolecules. Here, residue-selective deuteration of an FG Nup fragment is used to characterize its meso-scale structure and interactions with an NTR. The increased signal content compared to simpler SANS measurements requires enhanced signal processing and analysis. Results support the high degree of disorder present in apo- and in complex states with an NTR, the consistent presence of disorder of FG Nup chain complexes when interacting with multiple NTRs, and the free movement of FG motifs without significant inter-motif contacts.

## Results

### Scattering

SANS data was collected from five samples of [^2^H L-Phe]FSFG_12_ (17)(Table 1) spanning zero to two molar equivalents of NTR NTF2 (18) (Fig 1, left). As expected, these samples had about 10% of the scattering compared to perdeuterated similar materials(19). Even accounting for the relative error estimates, the scattering data exhibit sinusoidal additions to the normally smooth decays associated with intrinsically disordered proteins.

**Table 1.**

| ID | Composition<br>mM | D<sub>max</sub><br>GNOM | D<sub>max</sub><br>BIFT | R<sub>s</sub><sup>1</sup> GNOM | R<sub>s</sub> BIFT | R<sub>s</sub> Guinier |
| --- | --- | --- | --- | --- | --- | --- |
| A | FG<br>2.33 | 133 | 105 ± 6 | 43.0 ± 1.9 | 36.7 ± 1.4 | 39.1±1.2 |
| B | FG : NTF2<br>2.33:0.23 | 148 | 186 ± 1 | 44.6 ± 1.0 | 46.2 ± 0.8 | 42.2 ±1.3 |
| C | FG : NTF2<br>2.33:0.46 | 147 | 120 ± 3 | 44.1 ± 1.5 | 40.9 ± 0.3 | 42.6 ±1.3 |
| D | FG : NTF2<br>1.61:1.67 | 180 | 179 ± 6 | 57.8 ± 1.0 | 57.0 ± 0.3 | 56.3 ±1.9 |
| E | FG : NTF2<br>1.06:2.16 | 203 | 182 ± 4 | 63.9 ± 1.1 | 61.0 ± 0.4 | 58.6 ±1.7 |

**Figure 1.**
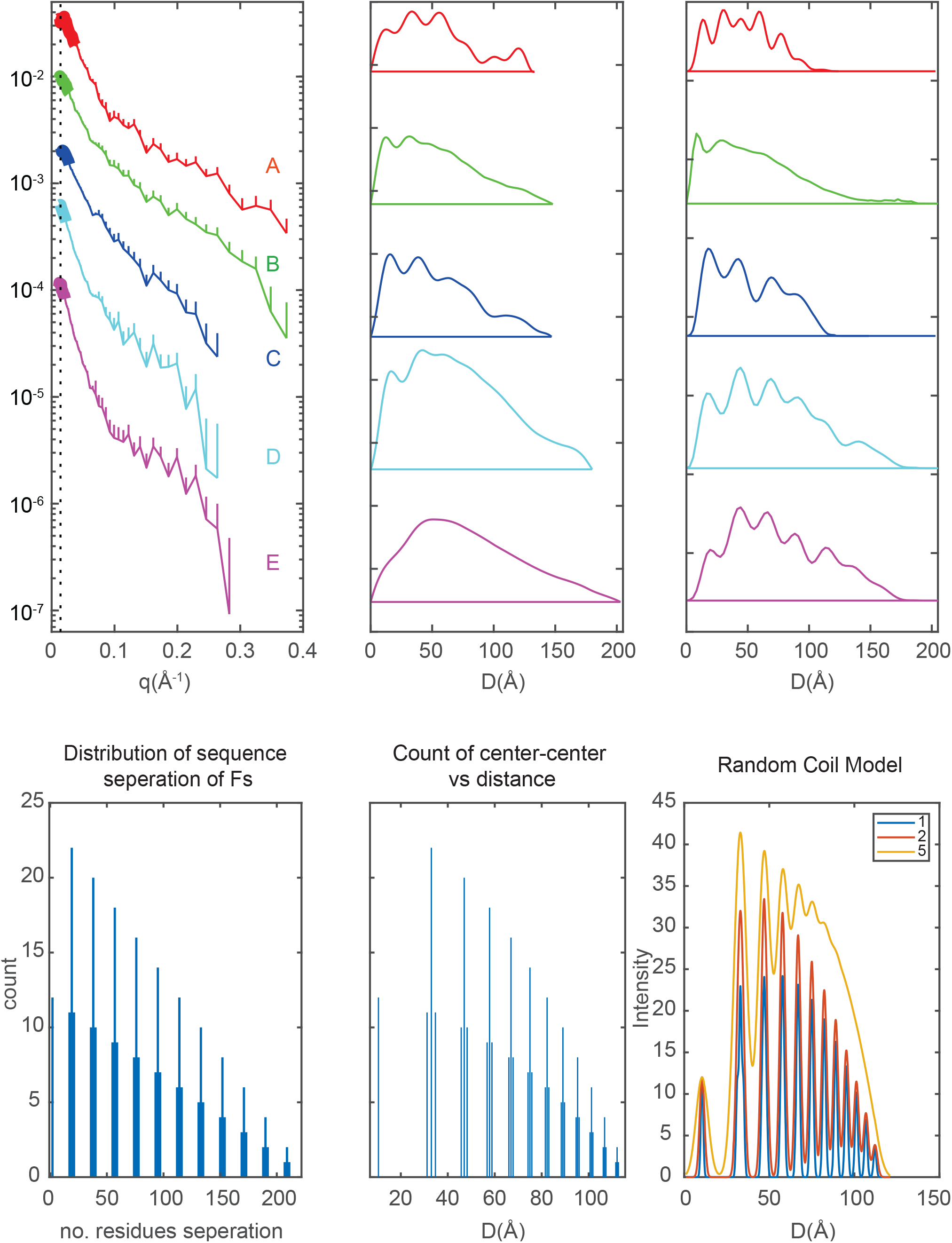
Observed scattering, PDDF transforms, and model-free predictions. (*Upper Left*) Observed scattering intensities of samples A-E (upper-lower) vs *q* (Å^-1^), with estimated error bars shown for upper range only. The black vertical at *q*=0.0141 is the lower limit of q used in *R*_*s*_^*1*^ calculations. The thickened regions are the regions selected in the Guinier analysis of *R*_*s*_ (*Center*) Probability distribution of scatterers for samples A-E vs separation distance *D*(Å) calculated by GNOM 5.0 (36), generally regarded as a stable converter of reciprocal and real distances with up to three oscillations (37). (*Right*) Bayesian Inverse FT Probability distribution of scatterers for samples A-E vs separation distance *D*(Å) calculated by BayesApp (38, 39) which uses a different approach to error analysis and minimizes arbitrary parameterization. *Bottom*: Random coil model prediction of (*left*) count of the separation of F-F residues pairs as a function of number of residues, *n*; (*center*) same ordinate data scaled to the expected distance for a random coil depending on *a*(*n*)^*ν*^ with values from (28) (*right*) previous panel arbitrarily Gaussian broadened with 1, 2, 5 Å half width.

### Inverse Fourier Transforms

Two slightly different inverse Fourier methods were applied to the original data (left panel, Fig 1) to obtain Paired Distance Distribution Functions (PDDFs). The widely used GNOM method involving parameters limiting the oscillations of the transform was applied, leading to the center panel (Fig 1), and the Bayesian Inverse Fourier Transform (BIFT) was used for the right panel. BIFT is designed to effectively assess over-, or underestimation of errors compared to experiments without fixed limits on oscillations. While the PDDFs have some similar characteristics, they clearly differ, with the BIFT set providing the appearance of finer structure. Analysis below will concentrate on the BIFT PDDFs.

### Modeling

We approached the derivation of structural details from the scattering PDDFs with two strategies (20, 21) -- (i) using atomic-model-free evaluation (AMF) by comparison of the PDDF with those predicted for standard polymer models, and (ii) model fitting of scattering to the predicted scattering of a pool of potential atomic scale models. These analyses assume a constant structure factor appropriate to dilute solution.

### Atomic-model-free: (AMF)

The inter-residue scattering from deuterons in a phenylalanyl side chain are at short distances and averaged by side chain rotations. A low-resolution model may then represent each F residue as a scattering center, a F-scatterer model. Given that FSFG_12_ is a random coil, the centers are then separated by the average dimensions of the sequence separating them. In Fig 1, lower left, the count of numbers of interactions as a function of sequence separation, *n*, is illustrated. This can then be rescaled on the abscissa as *r= a n*^*ν*^ where *a* is the average random coil separation, and *ν* is the coefficient from Flory theory (22) (Fig 1, lower center). Gaussian broadening then provides estimates of the effects of averaging (Fig 1, lower right). The experimental PDDFs do not conform directly to this F-scatterer model. The first peak in all PDDFs is at or greater than 13 Å .The *R*_*s*_ values observed (Table 1) and prior DLS data preclude significant intermolecular interactions of FSFG_12_, so this then represents the averaged interaction associated with the intramolecular **F**S**F** scattering and the average of all deuterons in each F residue. As the molar ratio of NTF2 is increased (Figs 1,2, from A to E) the distribution is more extended, although the distributions do not appear to be linear combinations of ‘free’ and ‘complexed’ sets, and some features associated with the highly random free material are still evident (e.g. 19, 44 Å on the E curve Fig 2). At the 1:2 molar ratio (which is close to full saturation (17)), the chain is significantly extended, and the regularity of F-F spacings is apparently modified, presumably by the very high number of possible conformers in the interacting forms. This arises from the twelve motifs of disordered FSFG_12_ interacting with the two sites on the NTF2 dimer, with 4-6 FSFG transiently interacting on one NTF2 site (17). When one NTF2 interacts with an FSFG_12,_ the space available for conformational averaging is reduced, so the apparent radius increases. The interaction of the second NTF2 results in a new pronounced feature at 113 Å reflecting the average site-site separation of FSFG interactions with two NTF2s as indicated by the stoichiometry of binding by ITC and NMR (17).

**2.**
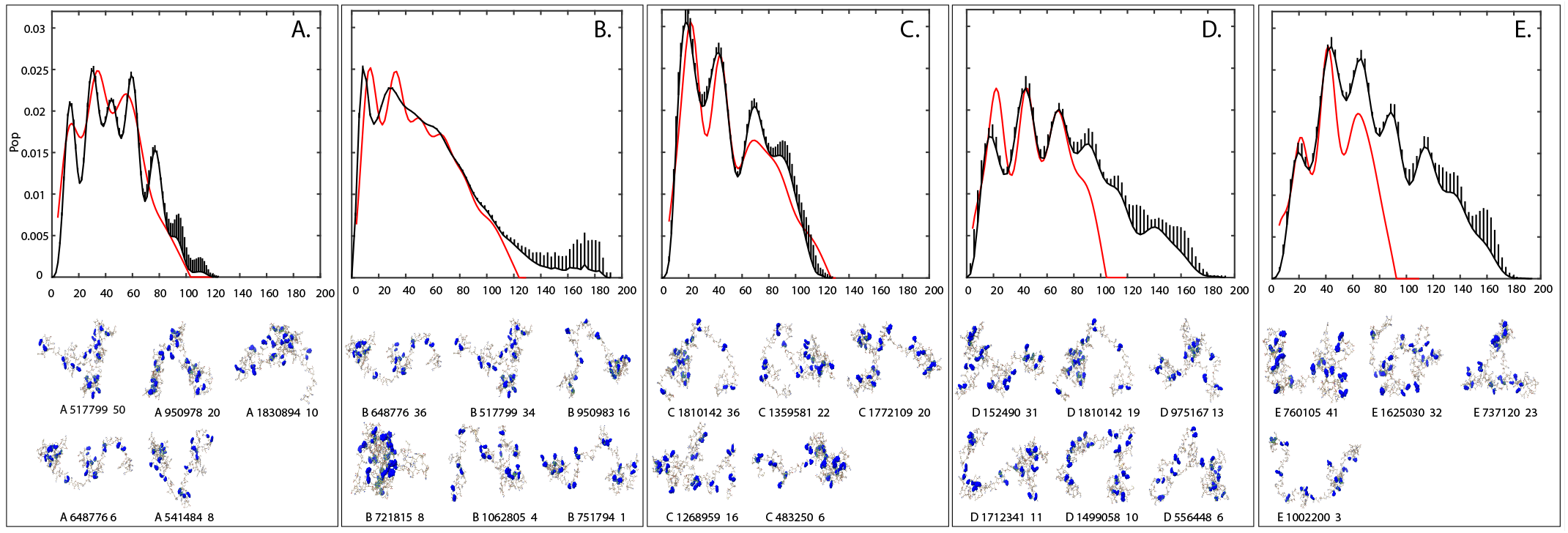
EOM Model fitting to scattering, illustrated by PDDFs. Ensemble of models approach (23, 40) was applied to fitting the scattering curves (Fig 1 right) for 2,000,006 models and results plotted in the PDDF space. For each panel, the BIFT PDDF is plotted in black with positive error bars (details in methods), and the fitted model summation is in red. (A) FSFG_12_ alone, (B) FSFG_12_ NTF2 (1:0.1 M:M), (C) FSFG_12_ NTF2, (1:0.2 M:M), (D) FSFG_12_ NTF2 (1:1 M:M), (E) FSFG_12_ NTF2 (1:2 M:M) (Table 1). The upper limits of estimated errors from BIFT are shown as vertical lines, at three times their actual values, for clarity. Note specifically that the fits calculated were to the observed scattering data, not the PDDFs. Inserted molecular figures below are the major contributors to the fitted curves, identified in the form <Panel number Structure number % contribution>, with Phe residues colored blue.

### Ensemble modeling

Using the Ensemble Optimization Method (EOM) (23) (Supp. Methods) for a set of random coil FSFG_12_ structures, the basic appearance of the PDDF is reproduced for traces A-C (Fig 1), while the set clearly does not contain extended structures similar to those expected for the complexes in traces D and E. At a low concentration of NTR, trace B, there is the appearance of a high degree of averaging, smearing the peak positions associated with the apo material (Fig 1. trace A) and the more complexed forms (C-E).

## Discussion

### FG Nup/ NTR interactions

The figures present simulations based on two separate model sets. The AMF (Fig 1 lower) system assumes a homogenous residue infinitely long polymer in a good solvent minimizing solvent interactions. The EOM (Fig 2) analysis uses an all-atom self-avoiding walk model, and is calibrated predominantly from folded structures and compared to random coil data (24). While neither consider specific interactions with solvent, nonetheless, the general features of the PDDFs and their correspondence to predicted (AFM) or fitted (EOM) systems allow detailed interpretation. Larger PDDF values do not correlate completely with the AFM series (Fig 1 lower) lacking its comb regularity, nor do they correlate completely with the EOM selections, but they do indicate spacing consistent with a high degree of random distribution, without any of the significant F/F interactions proposed previously for FG Nups e.g. (25). For the FSFG_12_ alone (A), the EOM fitting is marked by the lack of matching of number of peaks and their positions, suggesting that the generated pool lacks some features associated with this ensemble, or possibly that the BIFT has produced artifactual features. In contrast, panels B and C with 1:0.1 and 1:0.2 ratios of FSFG12:NTF2 display greater broadening of underlying features (compatible with the right lower AFM panel in Fig 1) reflecting a much larger pool of conformers in the rapid exchange with NTF2. The conformers fitted do not however appear to differ significantly from those in the apo material, indeed two (#517799 and #648776) are selected in both Fig 2 A and B. With higher ratios of NTF2, the fitting model ensemble lacks conformers directly reflecting NTF2 interactions and cannot match the scattering associated with greater than ∼ 100 Å separations (Fig 2, D, E). In Fig 2 E (1:2 molar ratio), the appearance of a peak at about 113 Å can be ascribed to an averaged separation of the FSFG interactions with two NTF2s. The detailed meso-scale scattering here is completely compatible with models of very rapid interactions between multiple sites of FG Nups and NTRs developed from solutions studies (17, 26, 27) and atomic MD simulation (28). They also exclude models of these interactions, at least for these classes of FG Nups and NTRs in similar concentration ranges, that include high degrees of interchain affinity and specific F-F interactions within FG Nups (29).

### Simulation development

Although the combination of SAS and MD calculations of complexes to compare predicted and observed data is appealing, there are two issues which discourage this here. First, while there has been significant development of force fields to describe IDP systems (e.g. (30)) there remains considerable controversy as to the appropriateness of current force fields (31). Indeed, given the unique meso-scale nature of the data here (10 - 200 Å)-- a range not used in force field optimization – an alternative objective may be adjustment of force fields to fit these and similar data. Second, the simulation of NTF2 binding to each of the FSFG motifs and in 1:1 and 2:1 stoichiometries for an estimated needed ∼10 μsec each (28) suggests a >500 μsec total simulation of a solvated multicomponent large box containing the intrinsically disordered systems, which is at the limit of current resources.

### SANS interpretive development : Prospective

This study demonstrates that site selective deuteration of proteins and solvent matching can provide unique information on the 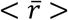 distribution of the scattering centers in multiple conditions not readily available from other methods. Improved measurement and analysis involving detector desmearing (32), additional detector positioning, and increased experience with the scattering-distance transform methods will improve precision of estimated distances. SANS has the specific advantage of broad applicability across all physical phases, so, e.g. structure factors can probe distributions within liquid-liquid phase separation (33) using dilution with unlabeled solutes to discriminate inter- and intra-molecular interactions. The general method is applicable also to carbohydrates, nucleic acids, and their complexes. While the current study probed the regular distribution of a single amino acid type, applications can employ multiple deuterated residue types, and multi-segmental synthesis (34). With increased resolution and sensitivity, the extension to the time domain of interaction using neutron spin echo and related methods (35) may be practical.

## Materials and Methods

### Summary of the main techniques

SI Appendix contains the detailed procedures for all the experiments and analysis. [^2^H] L-Phe was purchased from CIL Inc (Andover, MA) as 98.3% deuterated and > 98% chemical purity with no D-isomer detected.

### Protein expression and purification

The constructs used have previously been described (FSFG_12_, NTF2 -- (17)). To produce [^2^H L-Phe]-FSFG_12_, FSFG_12_ was recombinantly expressed in *E*.*coli* strain BL21(DE3) in minimal medium, and [^2^H] L-Phenylalanine 200 mg/L was added 30 mins before induction.

For SANS experiment, the stock solutions of [^2^H L-Phe] FSFG_12_ and NTF2 were dialyzed against 42% D_2_O buffer (20 mM HEPES, 150 mM KCl, 2 mM MgCl_2_, pH 6.8) in a common beaker. Samples A-E were prepared by adding the required volume of each protein from their respective stock solutions so as to make the final volume to 300 μL.

## Acknowledgments

We are grateful to the ORNL Bio-SANS group for their support and assistance. A portion of this research used resources at the High Flux Isotope Reactor, a DOE Office of Science User Facility operated by the Oak Ridge National Laboratory. Supported by grants from NIGMS R01 117212, 112108, DC is grateful to Charles Schwieters for help with Xplor-NIH.

## Supplementary Information for

### Preparation of materials

#### Protein expression and purification

Briefly, 5 μL glycerol stock of FSFG_12_-BL21 cells was revived by inoculating into 5 mL of LB medium and grown at 37 °C and 220 rpm for 8 h. The bacterial cells were pelleted down at 1,000 g for 10 m at room temperature and grown overnight in 100 mL M9 media containing ampicillin (100 μg/mL). From this, a 1 L M9 log-phase culture was prepared, and at an OD of 0.55 to 0.60, the culture was supplemented with 200 mg/L of [^2^H] L-phenylalanine and incubated for 30 m. The culture was induced with 1 mM IPTG (Gold Biotechnology) when the optical density at 600 nm reached 0.8 and was further grown for another 3 h. Induction was stopped by placing the culture on ice and cells were collected by centrifugation at 6,000 g for 20 m at 4 °C, washed with water, and stored at -80 °C.

For protein extraction, cell pellets were resuspended in the FG buffer (20 mM HEPES-KOH, pH 8.0, 150 mM KCl) containing 8 M urea supplemented with Pierce™ protease inhibitor mini tablets, EDTA-free (Thermo Scientific). After sonication, the lysate was ultracentrifuged at 40,000 g for 1 h at 4 °C, and the supernatant was filtered through a 0.45 μm filter. The [^2^H] L-Phe FSFG_12_ was purified using TALON resin (TaKaRa Bio USA, Inc.). Briefly, the resins were equilibrated with FG buffer with 8 M urea and incubated with the lysate for 20 m at room temperature. The column was washed with FG buffer containing 8 M urea, followed by 4 M urea, and further washed with FG buffer containing 2 mM imidazole. The [^2^H] L-Phe FSFG_12_ protein was eluted twice in elution buffers (20 mM HEPES-KOH pH 6.8, 150 mM KCl) containing 100 mM and 250 mM imidazole, respectively. The fractions were analyzed by 10% SDS-PAGE and the 100 mM imidazole eluted fraction was dialyzed against FG-buffer containing 2 mM MgCl_2_, pH 6.8, overnight. The dialyzed protein was concentrated using 3 kDa MWCO concentrators (Amicon) with dialysis buffer (20 mM HEPES-KOH pH 6.8, 150 mM KCl, 2 mM MgCl_2_, pH 6.8) containing 0.02 % NaN_3_.

To express NTF2 protein, pRSFDuet was utilized as the expression vector, and the transformed BL21DE3 cells (NEB) were cultured at 37°C in LB medium containing 50 μg/mL of kanamycin until they reached an OD600 of 0.8. Protein expression was induced by adding 1 mM IPTG to the culture and cells were harvested after 3 h. For purification, the cells were chemically lysed by using B-PER (Bacterial Protein Extraction Reagent) plus 20 mM MgCl_2_ according to the manufacturer’s instructions. Following a 15 m incubation period at room temperature with mild shaking, the suspension underwent an hour-long ultracentrifugation at 40,000 g at 4 °C. After the supernatant was incubated with TALON resins pre-equilibrated with NTF2 buffer (20 mM HEPES-NaOH pH 8.0, 150 mM NaCl) for 20 m at room temperature (gentle shaking), the unbound protein was removed by washing the column with NTF2 buffer and then NTF2 buffer containing 10 mM imidazole, pH 7.4. The NTF2 protein was eluted by NTF2 buffer (pH 7.4) containing 250 mM imidazole and the eluents were analyzed on a 12% SDS-PAGE. The fractions containing pure NTF2 were dialyzed against FG-buffer containing 2 mM MgCl_2_, pH 6.8, overnight.

The quantification of protein concentrations was done by BCA analysis (Pierce Co.), and proteins were stored at -20°C until ready to be used. Samples were stable for more than two weeks based on stable DOSY-NMR spectra. [^2^H] L-Phe incorporation into FSFG_12_ was assessed by comparison of integration of the aromatic ^1^H NMR region of the protein with unlabeled material and was 98.25% deuterated.

The sequence of the FSFG12 construct is

MNETSKPAFSFGAKSDEKKD DGDASKPAFSFGAKPDENKA ASATSKPAFSFGAKPEEKKD DDNSSKPAFSFGAKSNEDKQ QDGTAKPAFSFGAKPAEKNN NNETSKPAFSFGAKSDEKKD DGDASKPAFSFGAKSDEKKD DGDASKPAFSFGAKPDENKA ASATSKPAFSFGAKPEEKKD DDNSSKPAFSFGAKSNEDKQ QDGTAKPAFSFGAKPAEKNN NNETSKPAFSFGAKSDEKKD DGDASKPALEHHHHHH

### SANS data collection

For SANS experiment, the stock solutions of [^2^H L-Phe] FSFG_12_ and NTF2 were dialyzed against 42% D_2_O buffer (20 mM HEPES, 150 mM KCl, 2 mM MgCl_2_, pH 6.8) in a common beaker. Samples A-E were prepared by adding the required volume of each protein from their respective stock solutions so as to make the final volume up to 300 μL. Samples were stored frozen, thawed less than 6 h onto a 6°C cold stage, measured at 25°C. All samples remained clear with high viscosity.

Data was collected on the BioSANS HFIR ORNL instrument with standard conditions (19).

### Analysis of scattering data

Data reduction from SANS detectors used standard methods supplied by facility software drt-SANS for detector sensitivity correction, radical binning and background removal (19, 41). We constructed a two million ensemble of random structures to the FSFG_12_ sequence using TraDES (24) and added decoys of standard repeat peptide conformations and one ALPHAFOLD2 prediction. No decoys were ever detected in a fitting ensemble. GNOM or XPLOR-NIH (42) were used to derive SANS scattering. The EOM2.0 mode approach (40) was implemented in MATLAB to provide flexibility and speed using multiple cores. Two thousand chromosomes of 50-2,000 genes were mutated 20,000 times per step. Convergence based on stability of the derived *X*^*2*^ was achieved in less than 50 steps. However, this frequently reduced the number of contributing genes very significantly by duplication. An additional step beyond the EOM2.0 was introduced so that after detection of duplication an additional genetic algorithm was used to optimize the populations of the genes and the scattering baseline (16). This is similar to the optimization previously suggested (43) Details, illustrations of decoys and code are available at GitHub https://github.com/cowburn423/SANSE.

## Footnotes

1 In contrast to prior SANS studies, this approach reports on a radius of scattering *R*_*s*_, which will be smaller than a radius of gyration, because of the solvent matching of C- and N-termini and of any potential fixed water molecules.

